# Matrix metalloproteinase 1 modulates invasive behavior of tracheal branches during ingression into *Drosophila* flight muscles

**DOI:** 10.1101/669028

**Authors:** Julia Sauerwald, Wilko Backer, Till Matzat, Frank Schnorrer, Stefan Luschnig

## Abstract

Tubular networks like the vasculature extend branches throughout the bodies of animals, but how developing vessels interact with and invade tissues is not well understood. We investigated the underlying mechanisms using the developing tracheal tube network of *Drosophila* indirect flight muscles (IFMs) as a model. Live imaging revealed that tracheal sprouts invade IFMs directionally with growth-cone-like structures at branch tips. Ramification inside IFMs proceeds until tracheal branches fill the myotube. However, individual tracheal cells occupy largely separate territories, possibly mediated by cell-cell repulsion. Matrix metalloproteinase 1 (MMP1) is required in tracheal cells for normal invasion speed and for the dynamic organization of growth-cone-like branch tips. MMP1 remodels the Collagen IV-containing matrix around branch tips and promotes degradation of Branchless FGF in cultured cells. Thus, tracheal-derived MMP1 may play dual roles in sustaining branch invasion by modulating ECM properties as well as by shaping the distribution of the FGF chemoattractant.

## INTRODUCTION

Indirect flight muscles (IFMs) of flying insects display the highest known metabolic rates in the animal kingdom (Weis-Fogh, 1964). In Drosophila, two sets of IFMs, the dorsal-longitudinal muscles (DLMs) and the perpendicularly oriented dorso-ventral muscles (DVMs) are anchored to the thoracic cuticle and move the wings indirectly by deforming the thoracic exoskeleton rather than by acting directly on the wings. Each adult IFM is approximately 1 mm long and 100 µm wide (Spletter et al., 2018) and contains about 1000 nuclei (Rai & Nongthomba, 2013). To supply these large muscles with sufficient oxygen, an extensive network of gas-filled tracheal tubes not only superficially enwraps the IFMs, but also invades the myotube interior. This remarkable physiological adaptation minimizes the distance for oxygen diffusion from tracheoles to muscle mitochondria (Weis-Fogh, 1964; Wigglesworth & Lee, 1982) and provides efficient gas exchange for aerobic respiration to sustain flight over long time periods (Götz, 1987).

Tracheal cell migration is controlled by Fibroblast growth factor (FGF) signaling (Ghabrial et al., 2003; Hayashi & Kondo, 2018). The FGF ligand Branchless (Bnl) acts as a chemoattractant (Sutherland et al., 1996) that promotes tracheal cell motility by activating the receptor tyrosine-kinase (RTK) Breathless (Btl) on tracheal cells (Klämbt et al., 1992). IFMs receive their tracheal supply from tracheal cells that extend from the thoracic air sac primordium towards the notum region of the wing imaginal disc during larval development (Sato & Kornberg, 2002). Subsequently, during metamorphosis, tracheal terminal branches (tracheoles) ramify on and invade the developing IFMs (Peterson & Krasnow, 2015). Tracheal invasion into IFMs depends on the attraction of tracheal branches by Bnl FGF secreted on the muscle surface, followed by a switch to release of FGF from the interior transverse (T)-tubule system (Peterson & Krasnow, 2015). The T-tubule system is a network of tubular longitudinal and transversal membranes that extend around each sarcomere and are required for excitation-contraction coupling (Razzaq et al., 2001). It is continuous with the plasma membrane and was proposed to provide entry points for invasion of tracheal branches into the IFMs (Peterson & Krasnow, 2015). However, how tracheal cells interact with and enter the myotube, and how this process is coordinated with muscle development, is not clear.

Tracheal invasion into IFMs presumably requires dynamic remodeling of extracellular matrix (ECM) and plasma membranes, but the underlying mechanisms are not well understood. Matrix metalloproteinases (MMPs) are involved in tissue reorganization during branching morphogenesis in various systems, including the mammalian lung (Atkinson et al., 2005; Wiseman et al., 2003), mammary gland (Wiseman et al. 2003), and the *Drosophila* tracheal system (Page-McCaw et al., 2003). The *Drosophila* genome encodes two MMPs, MMP1 and MMP2, which perform common and distinct functions during tissue remodeling (Llano et al., 2002; Page-McCaw et al., 2003). MMP1 was shown to be required for tracheal remodeling during larval growth (Glasheen et al., 2009) and MMP2 for normal outgrowth of the air sac primordium (Wang et al., 2010). MMPs can be either secreted or membrane-tethered (LaFever et al., 2017; Page-McCaw et al., 2007), and are thought to function mainly as enzymes cleaving ECM components. However, MMP-mediated proteolysis can also modulate signaling by processing growth factors such as TNF*α*and TGF*β* (English et al., 2000; Yu & Stamenkovic, 2000), by regulating growth factor availability and mobility (S. Lee et al., 2005; Wang et al., 2010), or by cleaving growth factor receptors (Levi et al., 1996). MMP2 was shown to restrict FGF signaling through a lateral inhibition mechanism that maintains highest levels of FGF signaling in tracheal tip cells (Wang et al., 2010). Moreover, MMPs can regulate mammary gland development independently of their proteolytic activity (Kessenbrock et al., 2013; Mori et al., 2013).

To understand the mechanisms underlying tracheal invasion into IFMs, we analyzed the dynamics of the process *in vivo*. This revealed that tracheal cells invade IFMs directionally and migrate inside the myotubes with dynamic growth-cone-like structures at branch tips until tracheal branches fill the myotube volume. MMP1 activity is required in tracheal cells for normal invasive behavior and for the dynamic organization of growth-cone-like branch tips. We found that MMP1 remodels the Collagen IV-containing ECM surrounding invading branch tips and promotes degradation of Branchless FGF in cultured cells. Our results suggest that MMP1 activity may contribute to tracheal invasion not only by modulating local ECM properties, but also by altering the distribution of the FGF chemoattractant.

## RESULTS

### Tracheae invade flight muscles in a non-stereotyped, but coordinated manner

To understand the mode of IFM tracheation, we first analyzed tracheal branch pathways on and within IFMs. We focused our analysis on DLMs, which receive their tracheal supply from thoracic air sacs (Fig. 1A). Stochastic multicolor labeling of tracheal cells (Nern et al., 2015) revealed that multicellular air sacs converge into unicellular tubes (Fig. 1B) with ramified tracheal terminal cells at their ends (Fig. 1B’). Unlike tracheal terminal cells in other tissues, IFM tracheal cells not only ramify on the myotube surface, but also inside the syncytial myotube (Fig. 1C,C’ and D,D’; Supplementary Movie 1; Peterson & Krasnow, 2015). The cell bodies, including the nuclei, of IFM tracheal terminal cells reside on the myotube surface (Fig. 1C,C’’ and D,D’’), while IFM nuclei are distributed throughout the muscle between myofibril bundles as well as near the muscle surface (Fig. 1C,C’’’ and D,D’’’).

**Figure 1.**
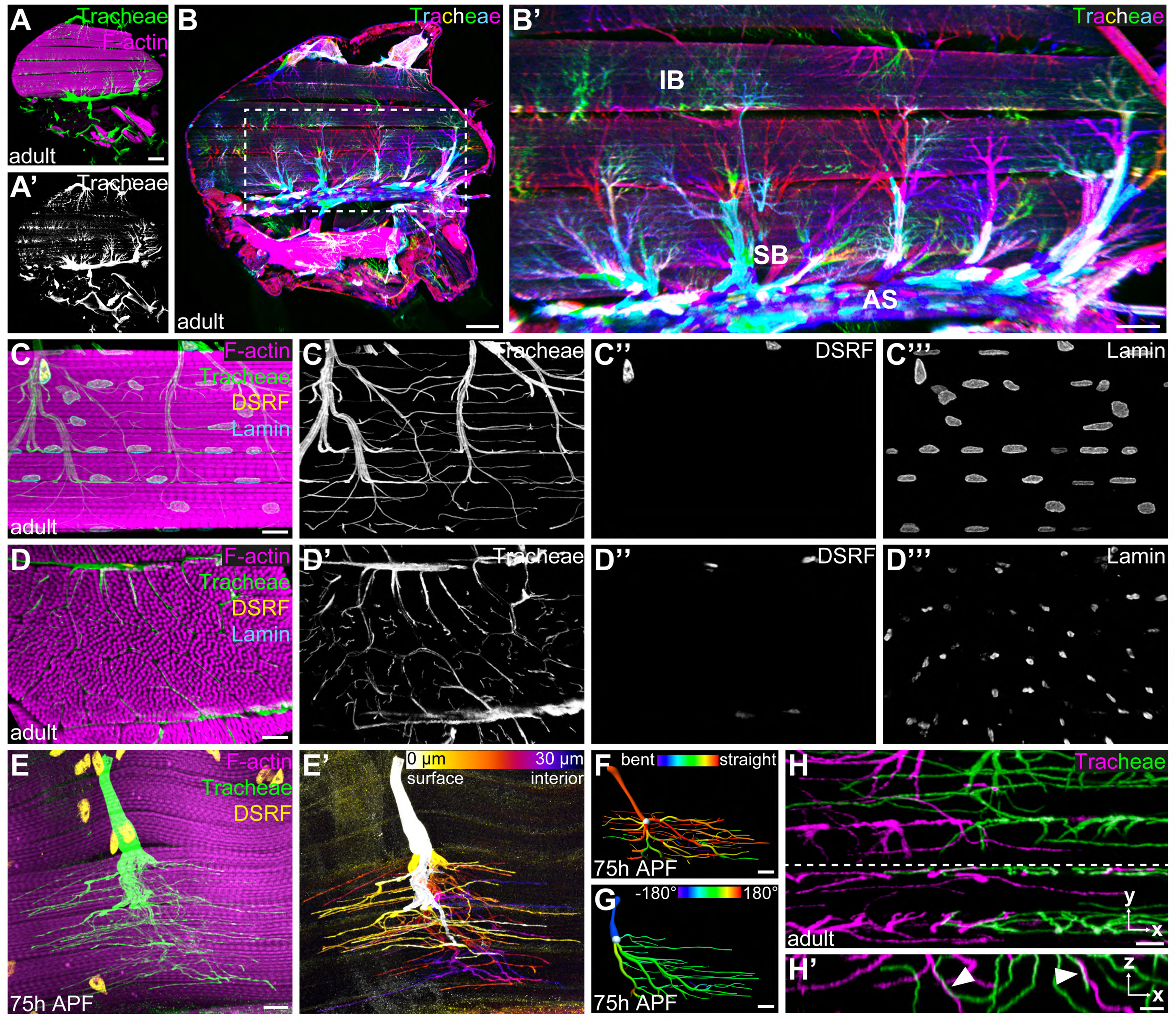
Tracheal terminal cell branches occupy separate territories in IFMs. **(A-A’)** Sagittal section of an adult thorax with dorsal longitudinal muscles (DLMs) stained for F-actin (magenta). Tracheal branches, visualized by their autofluorescence (green) arise from the thoracic air sacs adjacent to IFMs. (**B**) Stochastic multicolor labeling of tracheal cells in a sagittal section of an adult thorax. Multicellular tubes (**B’**) emanating from air sacs (AS) are superficial branches (SB) with tracheal terminal cells at their ends. Terminal cell branches spread on the muscle surface and invade as internal branches (IB) into the myotube. Note that individually labeled terminal cells occupy largely separate territories in IFMs (**B’**). **(C,D)** Tracheal branch supply of a single myotube in sagittal **(C-C’’’)** and cross-**(D-D’’’)** section stained for F-actin (magenta), LaminDm0 (cyan; all nuclei) and DSRF (yellow; tracheal terminal cell nuclei). Tracheal autofluorescence is shown in green. Cell bodies of tracheal terminal cells with DSRF-positive nuclei are located on the myotube surface. Terminal branches spread on the myotube surface, but also invade between and within myofibril bundles. **(E)** Single MARCM-labeled terminal tracheal cell in a developing IFM 75 h APF. The tracheal cell, labeled with cytoplasmic GFP (green) and nuclear DSRF (yellow), extends fine tracheoles parallel to myofibrils (magenta). Color-coding of depth (**E’**) indicates that a single tracheal cell ramifies deep into the myotube. The color map to the upper right indicates depth in the z-axis. **(F,G)** Segmented IFM tracheal terminal cell color-coded for branch straightness **(F)** and branch orientation angle **(G)**. Note that the majority of branches extend straight projections **(F)** and that branches extending from a given terminal cell often display a bias towards one orientation along the myotube long axis **(G). (H-H’)** Close-up of branches from two differentially labeled tracheal terminal cells (green and magenta). Note that while individual terminal cells occupy largely separate territories within the myotube, some branches of adjacent cells appear to be in close proximity or direct contact (arrowheads in **H’)**. H’ shows an orthogonal section in the x-z plane indicated by a dashed line in H. Scale bars: 100 μm (A, A’,B), 50 μm (B’), 10 μm (C-G), 5 μm (H,H’).

In each IFM, branch invasion starts between myofibril bundles and fine subcellular tracheal branches (tracheoles) also invade myofibril bundles (Fig. 1D,D’; Supplementary Movie 1), with most tracheoles extending parallel to the myofibrils (Fig. 1E,E’). The number and morphology of terminal cells supplying a specific DLM was variable between individuals, indicating that the tracheal branching pattern in IFMs is not stereotyped (Supplementary Fig. 1B). Interestingly, however, the number of tracheal branches was relatively uniform along DLM myotubes (Supplementary Fig. 1A, n=6). These findings suggest that branches originating from tracheal terminal cells uniformly fill the available myotube volume in a manner that is non-stereotyped, but tightly coordinated with myotube morphology.

### Tracheal cells occupy separate territories within the myotube

To investigate how tracheal cells arrange within myotubes to fill their volume, we generated animals carrying individually marked tracheal cells. Morphometric analysis of 31 individual IFM terminal tracheal cells revealed that these cells display highly variable cellular architectures, as measured by cellular volume, sum of branch lengths, and the number of branch and terminal points (Supplementary Fig. 1B, Supplementary Table 1). However, certain features were more uniform among cells. At least 95% of the branches of a given cell were aligned with the myotube axis (Fig. 1F; n=31) and the direction of branches was often biased towards one end of the myotube (Fig. 1G, Supplementary Fig. 1B). Stochastic multicolor labeling revealed that individual tracheal terminal cells occupy largely non-overlapping territories (Fig. 1B’). Interestingly, at the borders of such territories, branches from different cells were occasionally in close proximity or in direct contact (Fig. 1H,H’). These findings suggest that invading tracheal cells fill the available space within the myotube, but minimize overlaps, possibly mediated by contact-dependent repulsion between tracheal cells.

### Innervation precedes tracheal invasion into IFM myotubes

To investigate the dynamics of IFM tracheation, we analyzed a time course between 32 h APF and adulthood. While DVMs form de novo by fusion of adult muscle progenitors (AMPs), DLMs form by fusion of AMPs to larval ‘template’ muscles 8 h APF (Dutta et al., 2004; Fernandes et al., 1991). Tracheal invasion into DLMs begins around 48 h APF when tracheoles start to enwrap and invade the myotubes (Fig. 2A,A’; Peterson & Krasnow, 2015). Around 60 h APF DLM tracheation is not complete yet, indicating that tracheal ramification inside DLM myotubes continues during late pupal development (Fig. 2B,B’). Interestingly, prior to entry of tracheal branches into myotubes, motor neurons have already innervated IFMs (Fig. 2A,A’’). Furthermore, the distribution of tracheal branches was largely distinct from that of motor neurons (Fig. 2B,B’’) and the main tracheal branches did not overlap with motor neuron axons, suggesting that tracheae and neurons use separate entry routes into the muscle. Thus, tracheal invasion into myotubes is a comparatively slow process that occurs after IFM innervation.

**Figure 2.**
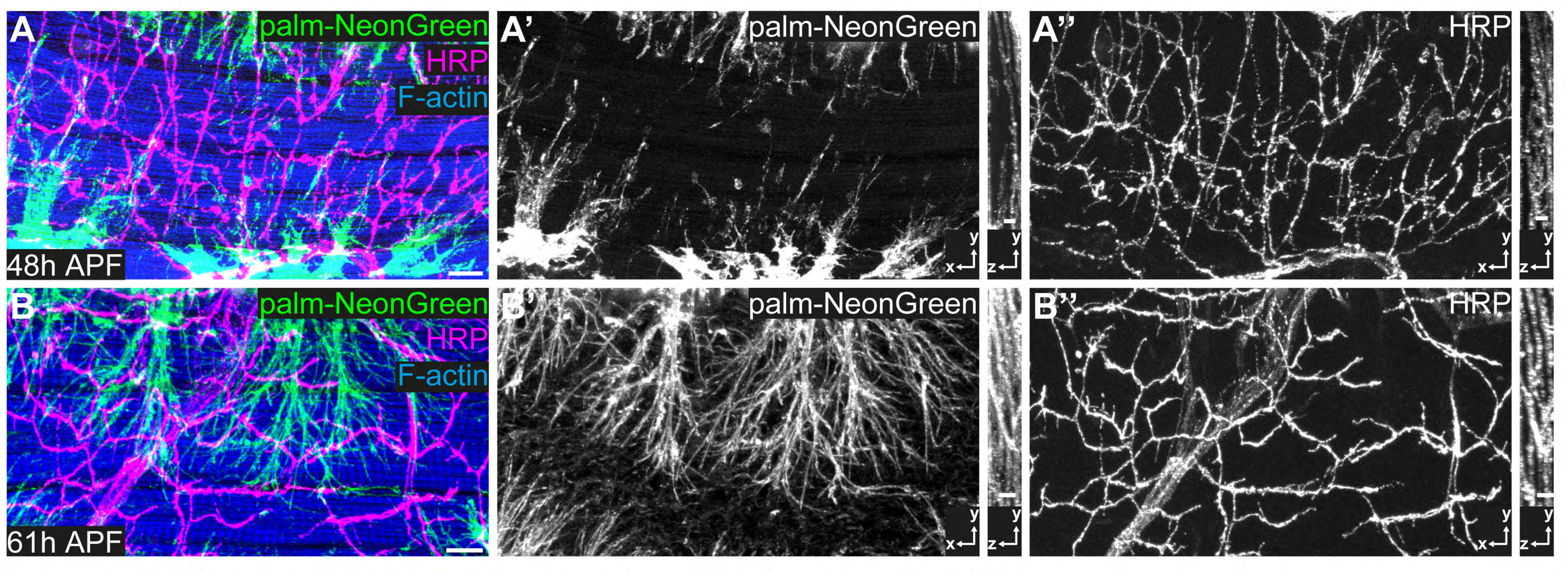
IFM innervation precedes tracheal invasion. **(A,B)** DLMs at 48 h APF (A) and 61 h APF (B). Tracheal cell membranes are labeled by palmitoylated mNeonGreen (green) driven by *btl*-Gal4. F-actin is labeled with Phalloidin (blue) and neurons with HRP (magenta). Note that at 48 h APF **(A-A’’)**, when tracheal invasion has just started, the muscle is already innervated with motor neurons. At 61 h APF tracheae have invaded the myotube (**B-B’’**). Cross-sections to the right show that most tracheal branches reside on the surface of the myotube at 48 h APF, whereas tracheal branches are inside the myotube at 61 h APF. Scale bars: 10 μm (A,B).

### IFM mitochondria enwrap tracheal cells, but not vice versa

Classical studies using dye infiltration experiments (Wigglesworth & Lee, 1982) described that IFM tracheole endings encircle IFM mitochondria, suggesting that mitochondria may be involved in guiding tracheal invasion inside the muscle. To investigate how mitochondria might influence tracheal branch pathways we analyzed the interrelationship of mitochondria and tracheae during IFM development using transmission electron microscopy (TEM). During sarcomere assembly in DLMs mitochondria change from a tubular morphology with few cristae to giant globular mitochondria with an elaborate cristae network (Supplementary Fig. 2A-F). Adult IFMs are packed with globular mitochondria between myofibrils, yielding tracheal branches closely associated with mitochondria along their entire length. However, in contrast to earlier reports (Wigglesworth & Lee, 1982), we were unable to detect any cases in which tracheal branches encircled mitochondria. Strikingly, however, we found that some mitochondria were partially enwrapped tracheal branches (Supplementary Fig. 2I-J; Supplementary Movie 2). These mitochondria showed no differences in volume or sphericity compared to mitochondria that were located farther away from tracheal branches (data not shown). Taken together, tracheal branches interact closely with mitochondria due to their dense packing between myofibrils and the partial enwrapping of tracheoles by mitochondria.

### *salm*-dependent flight muscle fate is required for tracheal invasion

We used tissue-specific RNAi to systematically search for tracheal- and muscle-derived factors, respectively, required for IFM invasion (Supplementary Fig. 3A). As previously reported (Peterson & Krasnow, 2015), the Bnl FGF chemoattractant is essential for IFM tracheation, as muscle-specific knock-down of Bnl completely abolished tracheal invasion into IFMs (Supplementary Fig. 3B,C). Interestingly, the trachealess muscles developed into adult IFMs with normal morphology of myofibrils, sarcomeres, mitochondria (Supplementary Fig. 3B,C,E,F), and with innervation by motor neurons (data not shown), suggesting that tracheal supply is dispensable for normal IFM development. However, adult flies lacking IFM tracheae were unable to fly. This finding prompted us to search for additional genes with roles in IFM tracheation using muscle-specific RNAi. We analyzed a set of 66 genes (Supplementary Table 2), which are required in IFMs for muscle function (flight), but not for normal muscle morphology (Schnorrer et al., 2010), suggesting that these genes may be involved in IFM tracheation. We used *Mef2*-Gal4 to knock down each of these genes in all muscles, and screened for changes in IFM tracheation. However, among the genes tested only the transcription factor Spalt major (Salm), which specifies fibrillar muscle fate (Schönbauer et al., 2011), was required for tracheal invasion into muscles (Supplementary Fig. 3D). Thus, as tracheal invasion only occurs in fibrillar muscles and not in other muscle types in *Drosophila* (Peterson & Krasnow, 2015), Salm-dependent processes appear to play a key role in preparing myotubes for tracheal invasion.

### MMP1 is required in tracheal cells for invasion into myotubes

We used an analogous RNAi approach to search for factors required in tracheal cells for branch invasion into myotubes (Supplementary Table 2) and identified an important role of matrix metalloproteinase 1 (MMP1) in this process. Knock-down of *Mmp1* in tracheal cells using *btl*-Gal4 led to altered tracheal branching on IFMs (Fig. 3A-C’). The angles between branches emanating from tracheal cell bodies on the myotube surface were reduced compared to control animals (Fig. 3E-G). In addition, fewer tracheal branches were found inside myotubes (Fig. 3H,I,J) and the fraction of myotube volume occupied by tracheoles was reduced (L). Furthermore, the tracheoles inside the myotubes showed fewer branch points compared to controls (Fig. 3M). Consistent with these findings, tracheal *Mmp1* knock-down led to reduced flight ability, indicating compromised muscle function of adult flies (Fig. 3N). We confirmed the specificity of the RNAi effect using two independent dsRNAs targeting different regions of the *Mmp1* gene, *Mmp1* RNAi 1 (JF01336, TRIP; Fig. 3B,B’,J,J’) and *Mmp1* RNAi 2 (Uhlirova & Bohmann, 2006), which led to comparable tracheal phenotypes (Fig. 3L,M). Furthermore, introducing the *Mmp1* homologue of *Drosophila pseudoobscura* under the control of its endogenous promoter (Ejsmont et al., 2009) into animals expressing *Mmp1* dsRNA in tracheal cells restored normal tracheal IFM invasion (Fig. 3C,C’,J,J’,L,M). Together these results indicate that the defects observed upon expression of *Mmp1* dsRNA are due to depletion of *Mmp1* (Fig. 3A-D).

**Figure 3.**
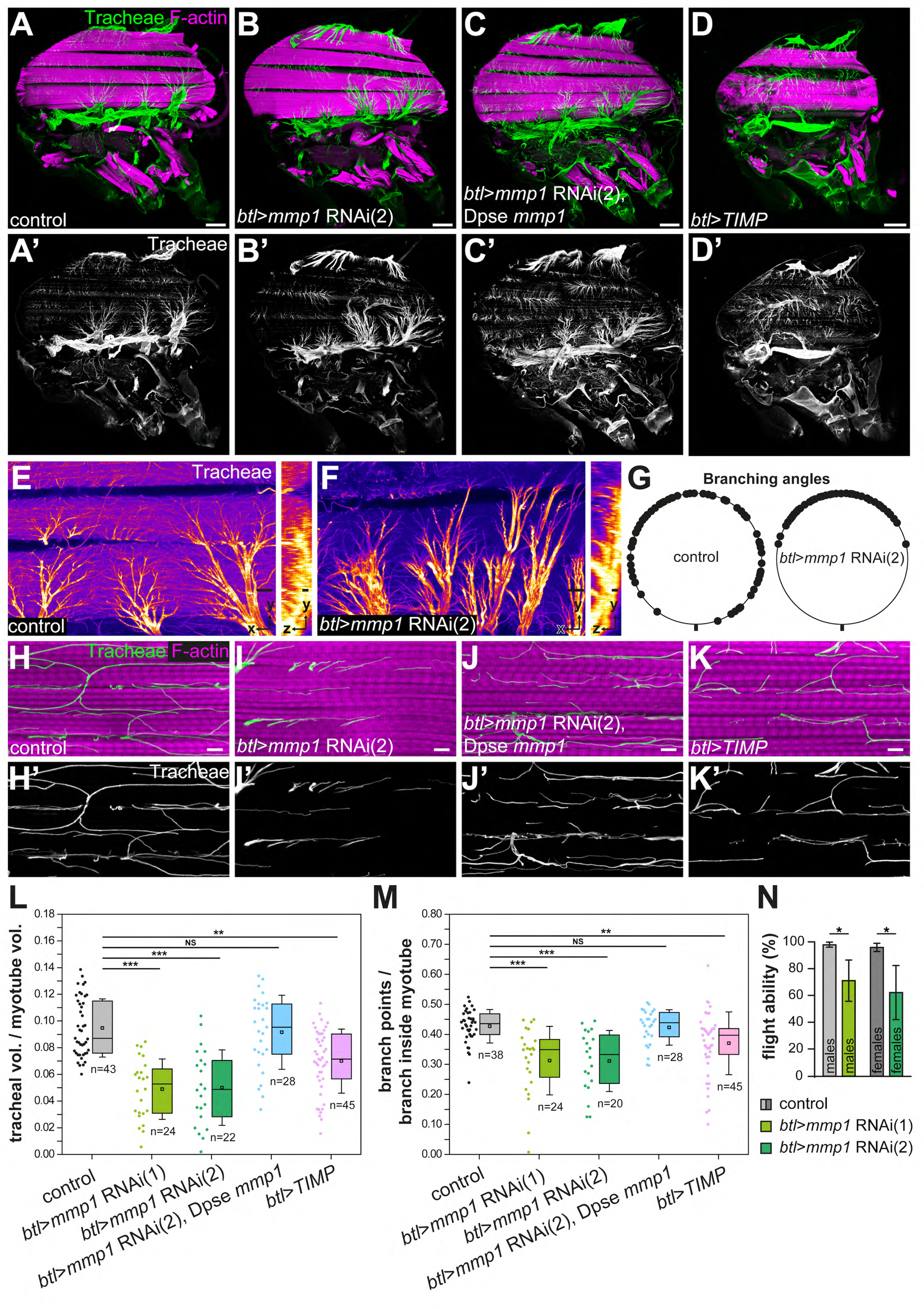
Tracheal branch invasion requires MMP1 function in tracheal cells. **(A-D’)** Sagittal sections of adult thoraxes stained for F-actin (magenta). Autofluorescence of tracheae is shown in green. **(E,F)** show close-ups of tracheal branches on the IFM surface. Orthogonal sections (yz) are shown to the right. Note altered spreading of tracheal branches on the myotube surface upon tracheal-specific *Mmp1* knock-down. The effect of *Mmp1* knock-down was rescued by the *Drosophila pseudoobscura Mmp1* homologue *GA18484* (Dpse *Mmp1*; **C**). Tracheal expression of TIMP (**D**) phenocopies the effect of RNAi-mediated *Mmp1* depletion. (**G**) The angles between branches emerging from tracheal cell bodies on the myotube surface are reduced in tracheal *Mmp1* knock-down animals compared to controls. (**H-K**) Sagittal sections of single myotubes. Note reduced number of tracheoles inside myotubes upon tracheal *Mmp1* knock-down (**I**) compared to control (**H**). The effect of *Mmp1* knock-down was rescued by the *Drosophila pseudoobscura Mmp1* homologue *GA18484* (Dpse *Mmp1*; **J**). Tracheal expression of TIMP (**K**) led to reduced tracheal invasion. (**L,M**) In a defined muscle volume the fraction occupied by tracheal branches (**L**) and the number of branch points per tracheal branch inside the myotube **(M)** were determined. At least 20 myotubes were scored. (**N**) Flight ability was measured as the percentage of flies that landed at the bottom immediately after discharging them into a Plexiglas cylinder (n=4 experiments). Note that tracheal *Mmp1* knock-down leads to reduced flight ability due to impaired muscle function. *Mmp1* was depleted using *Mmp1* RNAi (1) or *Mmp1* RNAi (2) (Uhlirova & Bohmann, 2006). * *P* < 0.05; ** *P* < 0.01; *** *P* < 0.001; NS not significant. Scale bars: 100 μm (A-D’), 20 μm (E,F) and 5 μm (H-K’).

Since Mmp1 has membrane-tethered and secreted isoforms (LaFever et al., 2017), Mmp1 could exhibit both cell-autonomous and cell-non-autonomous functions during branch invasion. To investigate Mmp1’s mode of action, we generated mosaic animals carrying clones of cells homozygous for the amorphic alleles *Mmp1*^*Q112*^ and *Mmp1*^*2*^ (Page-McCaw et al., 2003). However, we were not able to detect *Mmp1*^*Q112*^ or *Mmp1*^*2*^ mutant cells among IFM tracheae of mosaic animals (75 h APF). Although we cannot exclude the presence of additional cell-lethal mutations on the *Mmp1* mutant chromosomes, the absence of homozygous *Mmp1* mutant clones from IFMs suggests an essential cell-autonomous requirement of MMP1 in IFM tracheation. Together, these findings indicate that MMP1 is required in tracheal cells for normal invasion into IFMs.

### Tracheal invasion depends on MMP1 proteolytic activity

To test whether MMP1 catalytic activity, rather than a non-catalytic function (e.g. of the MMP1 hemopexin domains), was required for branch invasion, we expressed the *Drosophila* tissue inhibitor of metalloproteinases (TIMP; Pohar et al., 1999) in tracheal cells under the control of *btl*-Gal4. TIMPs inhibit MMP activity by occupying the active site of the protease (Gomis-Ruth et al., 1997), and *Drosophila* TIMP was shown to inhibit MMP1 and MMP2 (Page-McCaw et al., 2003; Wei et al., 2003). *btl*-Gal4-driven expression of TIMP resulted in air sac defects (Fig. 3D,D’), but adult flies were viable. The number of tracheal branches invading IFMs, as well as the number of tracheal branch points inside myotubes were reduced in these animals (Fig. 3K,K’,L,M), resembling the effect of tracheal *Mmp1* knock-down, although the defects caused by tracheal TIMP expression were milder compared to *Mmp1* knock-down. Together, these findings indicate that MMP1 catalytic activity is required in tracheal cells for normal IFM invasion.

### MMP1 modulates the speed of tracheal invasion into IFMs

To investigate the dynamics of IFM tracheal invasion and the function of MMP1 in this process, we developed a long-term live imaging protocol for visualizing IFM tracheation in living pupae (Fig. 4A). Tracheal cell membranes were labeled with *btl*-Gal4-driven palmitoylated mKate2 (palm-mKate2) and muscles were labeled with Myofilin-GFP. We imaged the onset of IFM tracheal invasion at 48 h APF, when tracheal branches extending from the air sac primordia begin to invade IFM myotubes (Fig. 4A,A’). Tracheal branches extending from the medioscutal air sac invaded the myotube in a directional fashion from posterior to anterior along the myotube long axis (Fig. 4B,B’ Supplementary Movie 3 upper panel). Such directional invasion of tracheae was also apparent for tracheal branches that extend from the lateroscutal air sac and invade dorso-ventral IFMs (Supplementary Movie 4 upper panel). Tracheal invasion upon tracheal-specific knock-down of *Mmp1* was still directional (Supplementary Movie 4 lower panel), but the extent of tracheal ramification inside the myotubes at 62 h APF was reduced compared to wild-type controls (Fig. 4 compare B,B’ to C,C’; Supplementary Movie 3). In control pupae the number of tracheal branches in a defined myotube volume close to the medioscutal air sac initially increased in a linear fashion and ceased at approximately 56 h APF (Fig. 4D; n=5). In contrast, *Mmp1*-depleted tracheal cells entered at constant, but lower speed into the myotube (Fig. 4D; n=8). Thus, tracheal MMP1 function is required for the normal dynamics and speed of IFM tracheation (Fig. 4D’).

**Figure 4.**
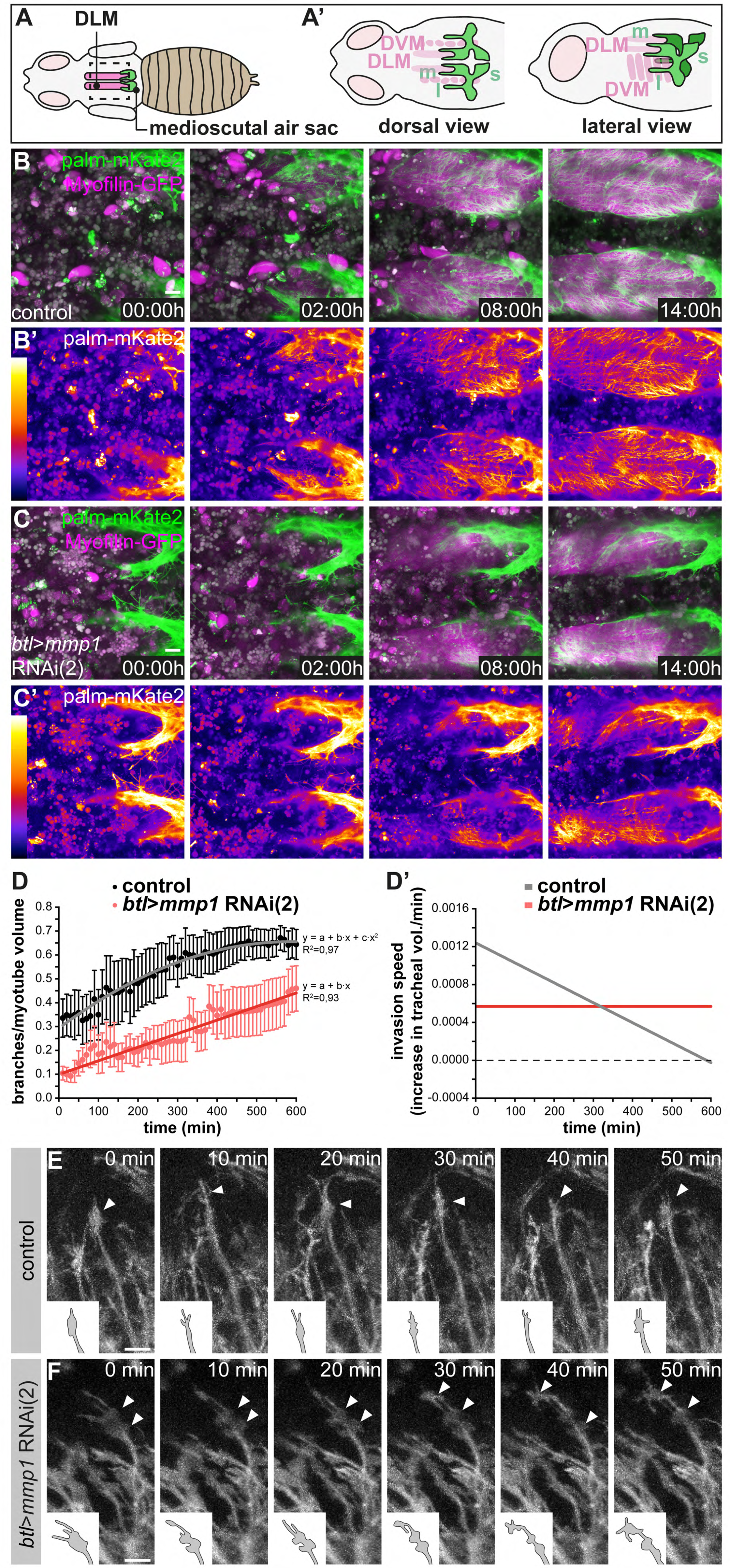
Normal dynamics of tracheal IFM invasion depends on tracheal Mmp1 function. (**A-A’**) Schematics of pupal flight muscles and air sacs 48 h APF. The pupal case around the head and thorax was removed for live imaging of tracheal invasion into IFMs. Tracheal invasion into dorsal longitudinal muscles (DLMs) from the medioscutal (m) air sacs was imaged from a dorsal view (**A’**). Tracheal invasion into dorso-ventral muscles (DVMs) from the lateroscutal (l) air sacs was imaged from a lateral view (**A’**). (**B,C**) Stills of tracheal invasion into DLMs by branches arising from the medioscutal air sac in a control pupa (**B,B’**) and a tracheal *Mmp1* knock-down pupa (**C,C’**). Palmitoylated mKate2 (palm-mKate2, green in B,C) labels tracheal cells, Myofilin-GFP (magenta in B,C) labels myotubes. B’ and C’ show palm-mKate2 intensities displayed as a heat map. The first time point (00:00h) corresponds to 48 h APF. (**D**) Quantification of tracheal branches over time in a defined myotube volume close to the medioscutal air sac. The speed of invasion (increase in tracheal branch fraction per minute; **D’**) was calculated for control and *Mmp1* knock-down pupae (n=3). (**E,F**) Stills of tracheal branch tips invading DVMs in control (**E**) and in tracheal *Mmp1* knock-down pupa (**F**). Note that growth-cone like structures (arrowheads) are confined to branch tips in the wild type, but are found also along branch stalks upon *Mmp1* knock-down (n=3). Scale bars: 20 μm (B,C) and 10 μm (E,F).

### Invading tracheae display growth cone-like structures at branch tips

The altered dynamics of branch invasion upon tracheal *Mmp1* knock-down suggests that MMP1 proteolysis could be required for clearance of the entry path for tracheae into the myotube or for modulating signaling molecules that promote branch invasion. To elucidate the role of MMP1 we investigated the behavior of individual tracheal branch tips in wild-type and in tracheal MMP1 knock-down animals. Invading branches displayed growth cone-like structures at their tips, with dynamic protrusions resembling lamellipodia and filopodia (Fig. 4E, Supplementary Movie 5). Growth-cone like structures at branch tips were also observed upon tracheal *Mmp1* knock-down (Fig. 4F, Supplementary Movie 5), and the presence of filopodia on these structures suggests that FGF signaling is active to promote migration also in cells with reduced MMP1 levels (Ribeiro et al., 2002). However, the growth-cone like structures in the MMP1-depleted cells did not remain confined to the branch tips as invasion proceeded (Fig. 4 compare E to F). Instead, multiple enlarged protrusions resembling lamellipodia persisted, often behind the branch tip, suggesting that trachea-derived MMP1 is required for the dynamic organization of growth cone-like structures during tracheal migration inside the myotube.

### Extracellular matrix components are distributed non-uniformly along invading tracheal branches

The reduced speed of tracheal branch invasion upon *Mmp1* knock-down could arise from a defect in the ability of tracheal cells to clear their path through the surrounding BM, the removal or remodeling of which may require MMP1 function. To investigate potential roles of MMP1 in BM remodeling during branch invasion, we first analyzed the distribution of the BM components CollagenIV-GFP, Laminin and Perlecan around tracheal branches at the onset of tracheal invasion 48 h APF. Compared to adult IFM tracheae, which were covered with Laminin- and Perlecan-containing BM (Supplementary Fig. 4C-C’’), invading tracheal branches at 48 h APF showed little detectable BM (Supplementary Fig. 4A-B’). The muscle surface, however, was covered with Laminin, Perlecan (Supplementary Fig. 4A-B’) and CollagenIV-GFP (data not shown). Of note, these ECM components also lined membrane invaginations on the myotube surface (Supplementary Fig. 4D and data not shown), presumably representing openings of the T-tubule network (Peterson & Krasnow, 2015). Entry of tracheal branches into these invaginations could require MMP activity. However, tracheal *Mmp1* knock-down did not notably affect the levels and distribution of Perlecan around branch tips that started to invade the myotube via the membrane invaginations (Supplementary Fig. 4E-F’).

Next, we analyzed the composition of the BM around tracheal branches in adult IFMs (Fig. 5; Supplementary Movie 6). In controls Laminin was detected along the entire length of tracheal branches, both on the muscle surface and inside myotubes (Fig. 5A,A’’,B,B’’; Supplementary Movie 6). In contrast, Perlecan and Collagen IV covered the tracheal stalks between myofibril bundles, but were not detectable around the tracheal tip regions inside myofibril bundles (Fig. 5A,A’,B,B’; Supplementary Movie 6), indicating that the BM around invading branch tips has a distinct composition and may be thinner than the BM surrounding the branch stalks.

**Figure 5.**
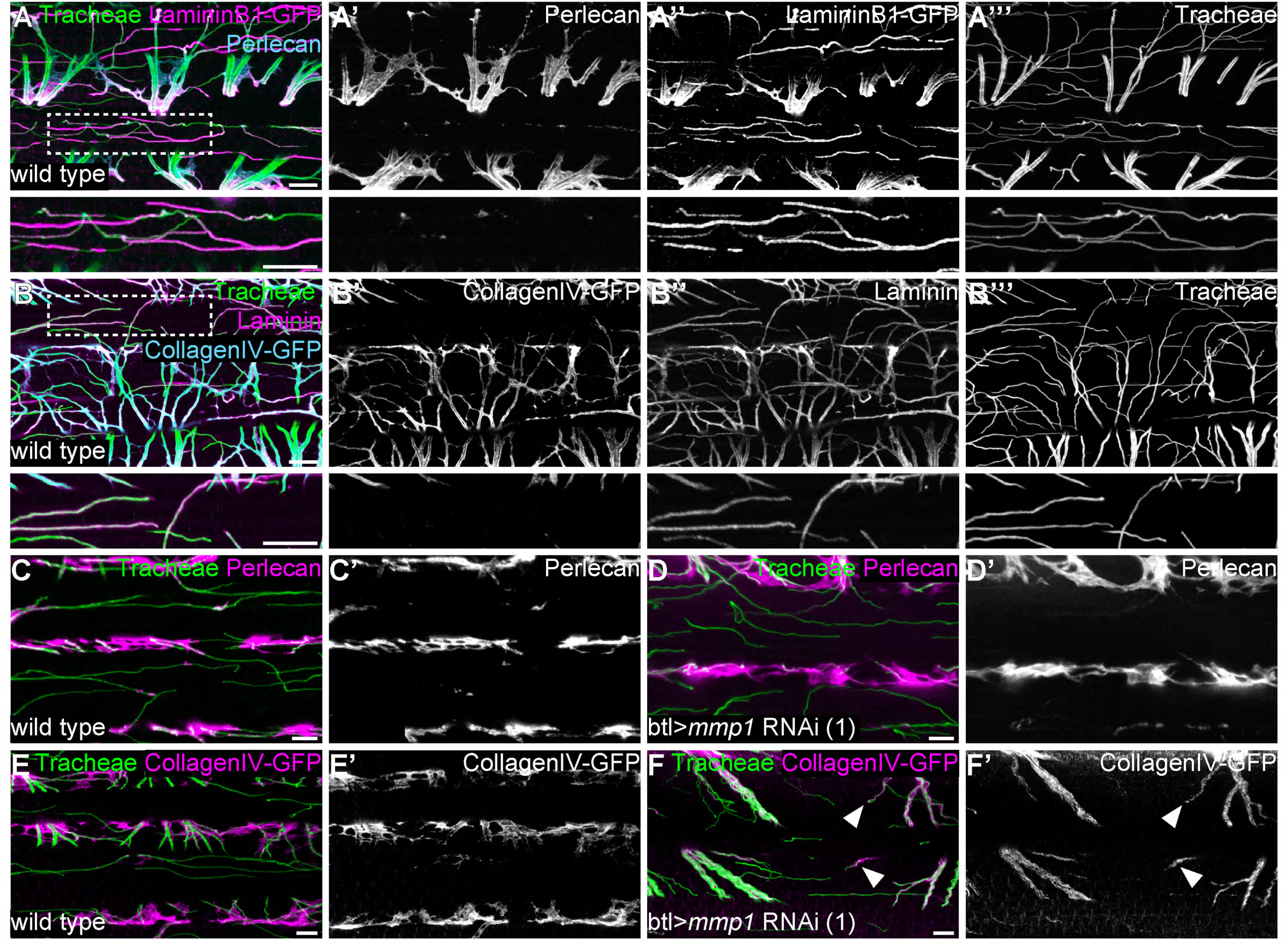
Tracheal stalk and tip regions show distinct basement membrane compositions. **(A,B)** Distribution of the ECM components Perlecan (**A,A’**), CollagenIV-GFP (**B,B’**) and Laminin or LamininB1-GFP (**A,A’’,B,B’’**) around tracheal branches inside an adult IFM myotube. Note that Laminin extends along the entire length of tracheal branches, whereas Perlecan and Collagen IV are excluded from branch tips inside myofibril bundles. Bottom panels show close-up view of the regions marked by dashed boxes. (**C-F**) Distribution of Perlecan (**C-D’**) and CollagenIV-GFP (**E-F’**) around tracheal branches in wild-type control (**C,C’,E,E’**) and *Mmp1* tracheal knock-down (**D,D’,F,F’**) adult myotubes. Note that Perlecan distribution is not affected by *Mmp1* knock-down, whereas CollagenIV-GFP extends into the tip region of some branches in *Mmp1* knock-down, but not in control animals (**F,F’** arrowheads). *Mmp1* was depleted using *Mmp1* RNAi (1). Scale bars: 7 μm (A-B’’’), 5 μm (C-F’).

### MMP1 is involved in remodeling of Collagen IV around tracheal branches inside IFM myotubes

We asked whether tracheal-derived MMP1 influences the distribution of BM membrane components associated with tracheal branches. Depletion of tracheal MMP1 did not appear to affect the levels and distribution of trachea-associated Perlecan (Fig. 5C-D’). However, *Mmp1* knock-down had a distinct effect on the distribution of Collagen IV. Whereas tracheal branch tips inside myotubes were devoid of Collagen IV in wild-type controls, 13% (n=59) of *Mmp1*-depleted tracheal branch tips showed Collagen IV signals around the tip region (Fig. 5E-F’). These findings suggest that trachea-derived MMP1 promotes invasion of tracheal branches through clearance of Collagen IV during migration inside the myotube.

### MMP1 promotes degradation of Bnl FGF in S2 cells

The mild effect of tracheal *Mmp1* knock-down on BM suggested that MMP1 activity might play additional roles independent of ECM remodeling. Given the established role of the chemoattractant Bnl FGF in tracheal cell migration, we asked whether MMP1 activity could influence Bnl FGF signaling. To test for effects of MMP1 on Bnl, we transfected *Drosophila* S2R^+^ cells with expression constructs for MMP1, Bnl FGF and the FGFR Breathless (Btl), and analyzed protein extracts from the cellular fraction and the culture medium (Fig. 6A). Bnl was detectable in protein extracts of the cellular fraction when Btl was co-expressed. Strikingly, co-expression of MMP1 led to a reduction of Bnl levels by on average 55% (n=3; Fig. 6A), suggesting that MMP1 activity can lead to degradation of Bnl protein. Due to the lack of tools to detect endogenous Bnl protein *in vivo*, we were unable to determine whether MMP1-mediated proteolysis plays a role in regulating the levels or distribution of Bnl FGF during IFM tracheation. However, these findings suggest the possibility that trachea-derived MMP1 activity could generate a local chemoattractant gradient by degrading IFM-derived Bnl FGF around the migrating tracheal branches.

**Figure 6.**
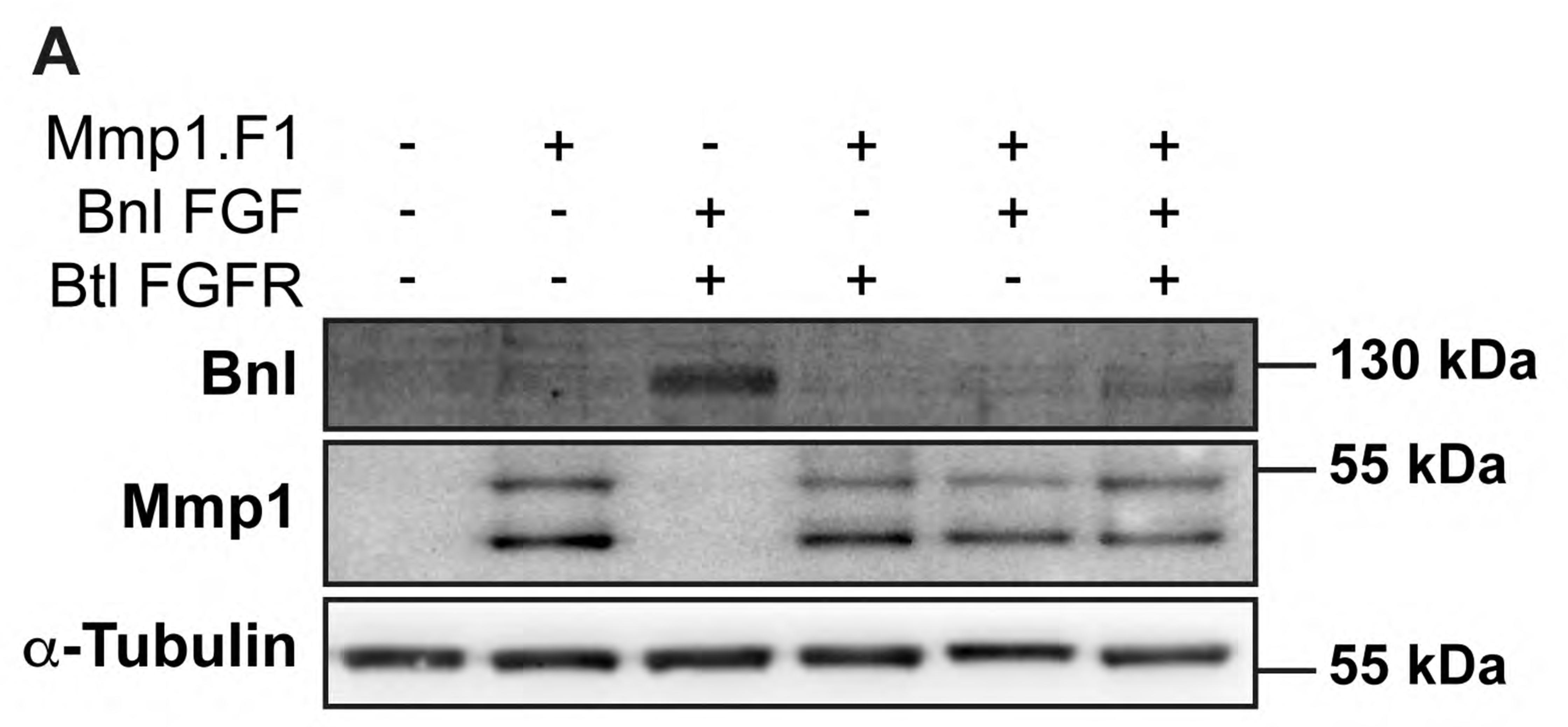
MMP1 promotes degradation of Bnl FGF in S2 cells. **(A)** Immunoblot of extracts from S2R^+^ cells probed with anti-Bnl (top panel), anti-MMP1 (middle panel), and anti-alpha-tubulin (lower panel) antibodies. Bnl FGF and Btl FGFR were expressed with or without MMP1 co-expression. Note that Bnl levels are reduced on average by 55% (n=3) in the presence of MMP1.

## DISCUSSION

Perfusion of tissues by oxygen-transporting vessels is a key prerequisite for all body functions in animals. In flying insects the extreme energy demands of flight are met by a network of tracheal tubes that minimize the distance for oxygen diffusion to mitochondria by extending branches into the interior of the flight muscles. This requires a new developmental process that enables tracheal cells to invade and spread throughout the IFM myotubes, unlike all other insect muscle types, where tracheoles ramify on the muscle surface only. Hence, flight muscle tracheation provides a powerful model to study the tissue interactions that promote branch invasion into tissues.

To investigate the cellular and molecular mechanisms underlying tracheal invasion into IFMs, we analyzed the dynamics of the process *in vivo*. First, through live imaging of muscle tracheation, we found that tracheal cells invade the muscle directionally with growth cone-like structures at branch tips. Tracheoles ramify inside the muscle until they uniformly fill the myotube volume. Intriguingly, however, single-cell analyses revealed that individual IFM tracheal cells occupy largely separate territories within the myotube, reminiscent of neuronal dendritic tiling (Grueber & Sagasti, 2010), suggesting that IFM tracheation involves repulsion between tracheal cells. Second, using a tissue-specific RNAi-based approach to identify factors required for branch invasion, we found that MMP1 activity is required in tracheal cells for normal speed of invasion and for the dynamic organization of growth-cone-like structures at migrating branch tips. Third, we showed that ECM components are distributed non-uniformly along IFM tracheal branches, with Laminin covering the entire length of tracheal branches, whereas Perlecan and Collagen IV are excluded from the tracheal tip regions inside the myotube. MMP1 is involved in remodeling the Collagen IV-containing matrix around invading branch tips. Interestingly, experiments in cultured cells revealed that MMP1 promotes degradation of Branchless FGF. Together, these findings suggest that MMP1 plays a dual role in sustaining tracheal branch invasion by remodeling the surrounding ECM as well as by modulating the distribution of the muscle-derived FGF chemoattractant.

A unique aspect of IFM tracheation is the fact that tracheal terminal cells enter and ramify within another cell, in this case a syncytial muscle. Although this system represents a specialized adaptation towards the extensive oxygen demand of this tissue, the underlying cellular mechanisms may be relevant also for the development of other organs such as the vasculature. Tracheal invasion involves dynamic adhesion to the substrate, guidance of tracheal cells, and remodeling of the myotube ECM and plasma membrane to accommodate the invading tubes. Angiogenesis, which is based on tip-cell-guided migration with invasive protrusions probing the environment, involves similar challenges, for instance in the case of blood vessels that grow into collagen-packed cornea tissue or into bones (Sivaraj & Adams, 2016).

The ability of tracheal cells to enter the IFM myotubes is likely to depend on permissive and instructive cues provided by the muscle, as well as on factors that act in the tracheal cells to mediate their invasive behavior. We showed that tracheal invasion into IFMs critically depends on the transcription factor Salm, which specifies the fibrillar muscle type (Schönbauer et al., 2011). The *salm*-dependent cell fate switch appears to induce a program that renders the myotube permissive for tracheal invasion, e.g. through modulating properties of the muscle plasma membrane or ECM. In addition, Salm may regulate factors that mediate the dynamic redistribution of the Bnl FGF chemoattractant from the muscle surface to the internal T-tubule network in IFMs. This switch in the mode of the subcellular pathway of FGF secretion was shown to guide tracheal cells into IFMs (Peterson & Krasnow, 2015).

Classical electron microscopy studies suggested that tracheoles enter the IFMs through plasma membrane invaginations that are continuous with T-tubules, and then spread through the T-tubule network (Smith, 1961a, 1961b; Wigglesworth & Lee, 1982). Other muscle types that lack these membrane invaginations are not invaded by tracheal branches (Peterson & Krasnow, 2015). Surprisingly, however, we did not find evidence that a normally organized T-tubule system is required for tracheal ingrowth and spreading in Drosophila IFMs, since *amphiphysin (amph)* mutants with a disorganized T-tubule system (Razzaq et al., 2001) showed a normal number and distribution of tracheoles in IFMs (Supplementary Fig. 4G,H and data not shown). Although the exact topology of the T-tubule system in wild-type and in *amph* mutant IFMs remain to be characterized, these results suggest that invasion into and spreading of tracheal cells inside IFMs does not depend on a pre-existing regular membrane invagination system.

IFM tracheoles are closely associated with mitochondria, thus minimizing the distance for gas exchange via diffusion. While we confirmed this close association by electron and high-resolution confocal microscopy, we found, contrary to an earlier report (Wigglesworth & Lee, 1982), no evidence that IFM tracheole endings encircle mitochondria. These earlier observations were based on dye infiltration experiments, which may lead to artifacts due to leakage of the injected dye used for tracheal staining. Conversely, we discovered mitochondria that were partially enwrapping IFM tracheoles. This is likely due to extensive fusion of mitochondria, resulting in giant sleeve-like mitochondrial geometries around tracheal tubes in IFMs. Intriguingly, this arrangement is reminiscent of the mitochondria that wrap around the axoneme in sperm tails (Woolley, 1970). Thus, mitochondrial wrapping may represent a common mechanism to sustain the extensive energy demands of specialized motile cell types such as flight muscle or sperm.

### MMP1 modulates invasive behavior of IFM tracheal cells

In addition to the muscle-derived factors discussed above, we show that the matrix metalloprotease MMP1 is required in IFM tracheal cells for their normal invasive behavior. ECM remodeling is crucial for branching morphogenesis of various organs, and MMPs are the main enzymes that mediate ECM degradation (Bonnans et al., 2014). We showed that while the BM around invading branches is very thin at the beginning of IFM tracheation at 48 h APF, tracheal branches in adult IFMs are surrounded by abundant, but molecularly heterogeneous, BM along their length. Laminin, but not Perlecan and Collagen IV, is present around branch tips. The presence of this molecularly distinct BM at the invasive branch tips suggests a reduced stiffness and increased distensibility of the BM, which has been observed also in other invading epithelia, including the mammalian salivary gland (Bernfield et al., 1972; Grobstein & Cohen, 1965; Harunaga et al., 2014), mammary gland (Fata et al., 2004) and lung (Moore et al., 2005). We found that depletion of MMP1 from tracheal cells led to a distinct effect on ECM remodeling. The tips of MMP1-depleted tracheal cells in mature IFMs showed residual Collagen IV, suggesting that MMP1 is involved in modulating the mechanical properties of the ECM surrounding invading tracheal branch tips by removing CollagenIV-containing ECM. The moderate strength of this effect in MMP1 knock-down animals may be attributable to incomplete RNAi-mediated depletion or genetic redundancy of MMP1 with other matrix-degrading proteases. Furthermore, we cannot rule out that MMP1 might act on other ECM components, besides Collagen IV, that we did not test.

Since MMP1 has membrane-tethered and secreted isoforms (LaFever et al., 2017), MMP1 may execute both cell-autonomous and cell-non-autonomous functions. However, our results based on genetic mosaic analysis suggest that MMP1 acts in a cell-autonomous manner during IFM invasion. This function relies largely on MMP proteolytic activity, as we showed by expressing the MMP activity inhibitor TIMP. These findings raise the question as to which are the relevant substrates of MMP1 activity.

### MMP1 might influence tracheal invasion by regulating FGF signaling

Historically, MMPs have been mainly associated with ECM remodeling (Bonnans et al., 2014). However, MMPS have broad substrate specificity and can cleave, besides several ECM components, also non-ECM proteins. Our finding that MMP1 promotes degradation of Bnl FGF in cultured cells suggests that MMP1 may be involved in regulating FGF signaling during IFM tracheal invasion. Directional migration of tracheal cells in the embryo is mediated by the graded distribution of the Bnl FGF chemoattractant, which is expressed locally in small clusters of cells (Sutherland et al., 1996). In case of the developing IFMs, the redirection of FGF secretion from the cell surface to T-tubules explains the switch from superficial to invasive tracheal cell migration (Peterson & Krasnow, 2015). Yet, it is not clear whether and how an FGF chemotactic gradient could be established along a syncytial muscle to control directional persistence of tracheal cell migration along the IFMs. Since MMP1 is expressed in the air sac primordium (Wang et al., 2010), tracheal cell-associated MMP1 might influence the spatial distribution of the FGF chemoattractant by degrading Bnl FGF and thereby generating a local sink of chemoattractant around the invading branches. Localized degradation of the FGF chemoattractant by the migrating branch tips could sustain motility of sprouting branches towards areas with higher concentrations of FGF, even if FGF is initially distributed uniformly on the muscle surface. Although we have not been able to visualize the distribution of endogenous Bnl FGF on pupal IFMs due to the limited sensitivity of the available tools, analogous chemotactic gradients generated by migrating cells have been described in different developmental processes (Dona et al., 2013; Tweedy et al., 2016; Venkiteswaran et al., 2013).

A regulatory interplay between MMPs and FGFs has been reported to operate also in other contexts of branching morphogenesis. MMP2-expressing tracheal tip cells are part of a lateral inhibition mechanism during larval *Drosophila* air sac development (Wang et al., 2010). Cells at the tip of the air sac primordium receive highest levels of FGF signaling and induce ERK-dependent gene expression, including induction of the *Mmp2* gene (Wang et al., 2010). MMP2 mediates release of an inhibitory signal that acts non-autonomously to prevent FGF signaling in neighboring cells and consequently restricts tip cell fate to the MMP2-expressing cells. The nature of the inhibitory signal is still unknown. Interestingly, expression of MMP2 can be induced by FGF2 in mammalian endothelial cells (Kohn et al., 1995). Mammalian MMP2 can also cleave FGFR1 to release a soluble receptor ectodomain fragment, which retains the ability to bind FGF and may influence FGF availability in the vascular BM (Levi et al., 1996). These findings suggest that MMP proteolytic activity may play a conserved role in modulating FGF signaling during branching morphogenesis in different developmental contexts across the animal kingdom.

## MATERIAL AND METHODS

### Drosophila strains and genetics

The following *Drosophila* stocks are described in FlyBase and were obtained from the Bloomington stock center, unless noted otherwise: *btl*-Gal4 (Shiga et al., 1996), *Mef2*-Gal4 (Ranganayakulu et al., 1995), UAS-*palm-mKate2* (Caviglia et al., 2016), UAS-*palm-mNeonGreen* (Sauerwald et al., 2017), UAS-*FRT>STOP>FRT-myr::smGFP-HA_V5_FLAG* (Nern et al., 2015), UAS-*Mmp1* RNAi (Uhlirova & Bohmann, 2006), *Mmp1*^*Q273*^, *Mmp1*^*Q112*^, *Mmp1*^*2*^(Glasheen et al., 2009), *amph*^*26*^ (Razzaq et al., 2001), Myofilin-GFP (fTRG501; Sarov et al., 2016; VDRC), LanB1-GFP (fTRG681; Sarov et al., 2016; VDRC), Vkg-GFP (G205; Buszczak et al., 2007). Additional UAS-RNAi stocks were obtained from the TRiP or VDRC collections (Supplementary Table 2). Crosses for RNAi knock-down were performed at 27°C. Heterozygous animals carrying the Gal4 driver (*Mef2*-Gal4 or *btl*-Gal4 crossed to *y w* flies) were used as controls in RNAi experiments. For rescue experiments, the fosmid clone FlyFos066598 (Ejsmont et al., 2009) containing the *Drosophila pseudoobscura Mmp1* homologue *GA18484* was integrated into the attP-9A landing site using PhiC31 integrase (Bischof et al., 2007).

### Genetic labeling of tracheal cell clones

For multicolor labeling of tracheal cells *btl-Gal4* flies were crossed to *hsFlp::Pest;; UAS-FRT>STOP>FRT-myr::smGFP-HA_V5_FLAG* (Nern et al., 2015). L3 larvae were heat-shocked for 20 min at 37°C. The MARCM system (T. Lee & Luo, 1999) was used for clonal labeling of tracheal cells. For wild-type MARCM clones *y w hs-Flp122; FRT40A tub-GAL80; btl-GAL4 UAS-GFP* females were crossed to *y w hs-Flp122; FRT40A FRTG13; btl-GAL4 UAS-GFP* males. For *Mmp1* mutant clones *y w hs-Flp122; FRTG13 tub-GAL80; btl-GAL4 UAS-GFP* females were crossed to males carrying *Mmp1*^*2*^, *Mmp1*^*Q112*^ or *Mmp1*^*Q273*^ mutations recombined on FRTG13 chromosomes. L3 larvae were heat-shocked for 2 h at 37°C. Pupae were staged by collecting white pre-pupae and dissected 75 h after puparium formation (APF) at 27°C.

### Flight test

Adult flies (one to five days after eclosion) were kept for four days at 30°C. before testing for flight ability in a Plexiglas cylinder as described in Weitkunat and Schnorrer (2014). We determined the percentage of flies that landed at the bottom immediately after discharging them into the Plexiglas cylinder.

### Cell culture and transfections

Plasmids used in transfection experiments were *Act5C-Gal4* (gift from Sven Bogdan, University of Marburg, Germany), *pUAST-Mmp1.F1* (Page-McCaw et al., 2003), *pMK33-Bnl-FLAG-HA* (DGRC), *pUAST-btl* (gift from Markus Affolter, University of Basel, Switzerland) and *pUAST-attB-K7*. The latter is a derivative of *pUAST-attB* (Bischof et al., 2007), in which a deletion of the SV40 3’-UTR results in higher expression levels due to loss of sensitivity to nonsense-mediated mRNA decay (Nelson et al., 2018). *Drosophila* S2R^+^cells (Yanagawa et al., 1998) were propagated in Schneider’s *Drosophila* medium (Gibco) supplemented with 10% FBS, 50 units/mL penicillin and 50 μg/mL streptomycin in 75 cm^2^ T-flasks (Sarstedt) at 25°C. Plasmids were transfected using Fugene reagent (Promega). *Act5C-Gal4* DNA (0.6 μg), *pUAST* and *pMT* constructs (in total 1 µg) were incubated with Fugene reagent for 30 min and added dropwise to the cells, which were seeded in 24-well plates on the previous day (3×10^5^ cells per well). Transfection medium was replaced by fresh medium without FBS and antibiotics 48 h after transfection. Metallothionein promoter-driven gene expression was induced by adding CuSO_4_ to a final concentration of 500 μM for 24 h.

### Immunoblots

Cells were harvested and lysed in ice-cold lysis buffer (20 mM Tris pH 8, 200 mM NaCl, 1 mM EDTA, 0.5% NP40) with protease inhibitor cocktail (Roche). Protein content of extracts was estimated using Pierce 660 nm Protein Assay Reagent (Thermo Fisher) supplemented with Ionic Detergent Compatibility Reagent (Thermo Fisher) for samples in Lämmli buffer.

Antibodies were mouse anti-Tubulin (1:1,000; Sigma DM1A), rabbit anti-Bnl (1:100; Jarecki et al., 1999; Sutherland et al., 1996), mixture of anti-MMP1 3B8, 3A6, 5H7 (1:10/1:100/1:100; DSHB; Page-McCaw et al., 2003), goat anti-rabbit Superclonal HRP conjugate (1:3,000; Thermo Fisher) and goat anti-mouse Superclonal HRP conjugate (1:3,000; Thermo Fisher). Protein extracts were separated on 12.5% SDS polyacrylamide gels (35 μg protein per lane) and electro-transferred to PVDF membranes. Bound secondary antibodies were visualized using the ECL-system (Amersham). Three independent samples were analyzed. Quantification of Western blot bands was performed using GelAnalyzer2010a with background subtraction.

### Immunostainings

For immunostainings of developing IFMs, white pre-pupae were staged at 27°C and dissected at the desired time APF. Pupae up to 60 h APF were dissected according to the protocol for developing IFMs (Weitkunat & Schnorrer, 2014). Older pupae and adults (1-6 days after eclosion) were dissected according to the protocol for adult IFMs (Weitkunat & Schnorrer, 2014).

Primary antibodies were chicken anti-GFP (1:500; Abcam ab13970), mouse anti-DSRF (1:100; Samakovlis et al., 1996), chicken anti-HA (1:200; Abcam ab9111), anti-FLAG(M2) (1:1000; Sigma F1804), mouse anti-Dlg1 (1:200; DSHB 4F3), guinea-pig anti-LaminDm0 (1:1000; gift from Georg Krohne, University of Würzburg), rabbit anti-Laminin (1:1500; Schneider et al., 2006), rabbit anti-Perlecan (1:2000; Schneider et al., 2006), anti-ATP5A (1:100; Abcam ab14748). Goat secondary antibodies were conjugated with Alexa Fluor 488/Dylight 488 (1:500; Life Technologies), Alexa Fluor 555/568 (1:500; Life Technologies), Dylight 550 (1:500; Thermo Fisher) or Alexa Fluor 647 (1:500; Life Technologies). Tracheae in adult IFMs were visualized by their autofluorescence upon UV (405 nm) excitation. Phalloidin-TRITC (1:1000; Sigma) was used to stain F-actin and anti-HRP-Alexa Fluor 647 (1:100; Dianova) to label neurons.

### Transmission electron microscopy

Pupal and adult IFM samples were fixed at room temperature (RT) in 4% paraformaldehyde and 0.5% glutaraldehyde in 0.1 M phosphate buffer (PB) for at least 2 h and were then transferred to 4 °C overnight. On the next day, samples were fixed with 2% OsO_4_ in 0.1 M PB for 1 h on ice in the dark. Next, samples were washed five times using ddH_2_O and stained *en bloc* with 2% uranyl acetate (UA) for 30 min at RT in the dark. After five washes in ddH_2_O, samples were dehydrated in an ethanol series (50%, 70%, 80%, 90%, 95%), each for 3 min on ice and twice in 100% ethanol for 5 min at RT. Following dehydration, samples were incubated twice in pure propylene oxide (PO) for 15 min and then transferred to an epon-PO mixture (1:1) to allow resin penetration over night. After removal of PO by slow evaporation over 24 h, samples were embedded in freshly prepared epon (polymerization at 60°C for 24 h). 90 nm sections were prepared on a Leica UC7 microtome and stained with 2% UA for 30 min and 0.4% lead citrate for 3 min to enhance contrast. Images were acquired with a Zeiss EM900 (80 kV) using a side-mounted camera (Osis).

### Light microscopy and image analysis

Stained specimens were imaged on a Zeiss LSM880 Airyscan confocal microscope or a Zeiss LSM710 confocal microscope. Z-projections were generated with Imaris (v9, Bitplane) using the ‘3D view’ or with Fiji (GPL v2; Schindelin et al., 2012). Where indicated, images were acquired with the Airyscan detector and subjected to Airyscan processing.

### Live imaging of tracheal invasion into pupal IFMs

Live imaging of IFM tracheation was performed on a Leica SP8 confocal microscope using a 40x/1.3 NA oil immersion objective, resonant scanning mode and Hybrid Detectors. Pupae expressing *btl*-Gal4-driven palmitoylated mKate2 (palm-mKate2; Caviglia et al., 2016) to label tracheal cells and Myofilin-GFP (fTRG501; Sarov et al., 2016) to label muscles were prepared for live imaging 48 h APF after staging at 27°C. The pupal case around the head and thorax was removed using forceps. Pupae were fixed on a coverslip with heptane glue and covered with a gas-permeable membrane (bioFOLIE 25, In Vitro System and Services, Göttingen, Germany) using a spacer of 0.5 mm. The dorsal-most DLMs were imaged from a dorsal view. DVMs were imaged from a lateral view. Time-lapse movies were recorded with z-stacks (100 μm thickness, 0.35 μm step size) acquired every 10 min over 14 h.

### Analysis of tracheal invasion speed

To quantify the progress of IFM tracheation over time movies were processed with Fiji. A binary movie sequence of tracheal invasion was generated by first creating a maximum intensity projection, followed by histogram normalization (1% saturation) and auto thresholding (Huang filter). In a ROI of approximately 1680 µm^2^ close to the medioscutal air sacs, changes in tracheal area fraction starting from 52 h APF were assessed over time with the Time Series Analyzer Plugin.

### Quantification of tracheal density in IFMs

Confocal sections of tracheae in IFMs were taken below the muscle surface. Stacks of 9 μm thickness (step size 0.3 μm) were acquired along the selected myotube in different individuals. Tracheal branches were visualized in a myotube volume of approximately 3×10^4^µm^3^. After average intensity projection, background subtraction (sliding paraboloid, radius 15), median filtering (radius 2), histogram normalization (1% saturation) and manual thresholding was performed in Fiji to generate binary images of tracheal branches. These binary images were used to determine the fraction of myotube area occupied by tracheal branches.

### Quantification of tracheal branch points per myotube volume

To determine the number of tracheal branch points in a myotube volume of approximately 3×10^4^ µm^3^ the same image raw data as described above were used. 3D binary images of tracheal branches were generated in Fiji by background subtraction (sliding paraboloid, radius 15), median filtering (radius 2), histogram normalization (1% saturation), Gaussian blur (sigma 0.2) and manual thresholding. The 3D binary images were subjected to the Skeletonize 3D plugin of Fiji to count the number of branches and branch points.

### Analysis of tracheal branching angles

The angles of branches extending from tracheal cell bodies on the IFM surface were determined by superimposing a circle with defined radius over a tracheal branch tree such that the circle covers the branch tree and the stalk. The intersections of tracheal branches with the circumference of the circle were recorded and plotted.

### Morphometry of tracheal cells

Z-stacks of single MARCM-labeled tracheal cells were acquired with a step size of 0.5 μm. Brightness correction along z was applied during image acquisition to visualize finest branches deep inside the myotube. 3D binary images of tracheal branches were generated in Fiji by background subtraction (sliding paraboloid, radius 15), median filtering (radius 2) and manual thresholding, and were segmented in Imaris. The Surface tool of Imaris was used to determine the cellular volume. The Filament tracer tool was used to segment branches and to extract the sum of branch length, total number of branch points and terminal points, branch straightness, branch orientation angle and branching angle. Automated filament tracing was adjusted manually for correctness.

### Analysis of mitochondrial morphology

Airyscan images of IFM mitochondria were acquired with a step size of 0.3 μm. Mitochondria were segmented using the Surface tool of Imaris with “split touching objects” enabled. Analysis of segmented mitochondria was performed using the Vantage tool of Imaris.

### Statistics and reproducibility

For phenotypic analyses, sample size (n) was not predetermined using statistical methods, but was assessed by taking into account the variability of a given phenotype, determined by the standard deviation. Experiments were considered independent if the specimens analyzed were derived from different parental crosses. During experiments investigators were not blinded to allocation. For statistical analysis, the Kolmogorov-Smirnov test was applied, which does not require assumptions on the type of data distribution.

## ACKNOWLEDEGMENTS

We thank Anna Körte for help with the RNAi screen, Aynur Kaya-Çopur for introducing us to flight muscle dissection and for providing samples for electron microscopy analysis, and Manuel Hollmann and Mylène Lancino for comments on the manuscript. We thank Markus Affolter, Stefan Baumgartner, Andrea Page-McCaw and Mirka Uhlirova for providing fly stocks and reagents. We are grateful to Christian Lehner for discussions and support at the Institute of Molecular Life Sciences (University of Zurich) during the first phase of this project. We are grateful to Christian Klämbt for discussions and for providing access to facilities at the Institute of Neuro- and Behavioral Biology (University of Münster).

## AUTHOR CONTRIBUTIONS

Conceptualization: JS, FS, SL

Data curation: JS

Formal analysis: JS

Funding acquisition: JS, SL

Investigation: JS, WB, TM

Methodology: JS, WB, TM

Supervision: SL

Visualization: JS, TM

Writing - original draft: JS

Writing - review & editing: JS, FS, SL

Unless noted otherwise the experiments were carried out by JS with the help of WB. Electron microscopy experiments were conducted by TM. Experimental work was planned by JS with SL and FS. Data were analyzed by JS. The manuscript was written by JS and SL with feedback by FS. SL conceived and supervised the study with input from FS.

## Supplementary Figures

**Supplementary Figure 1.**
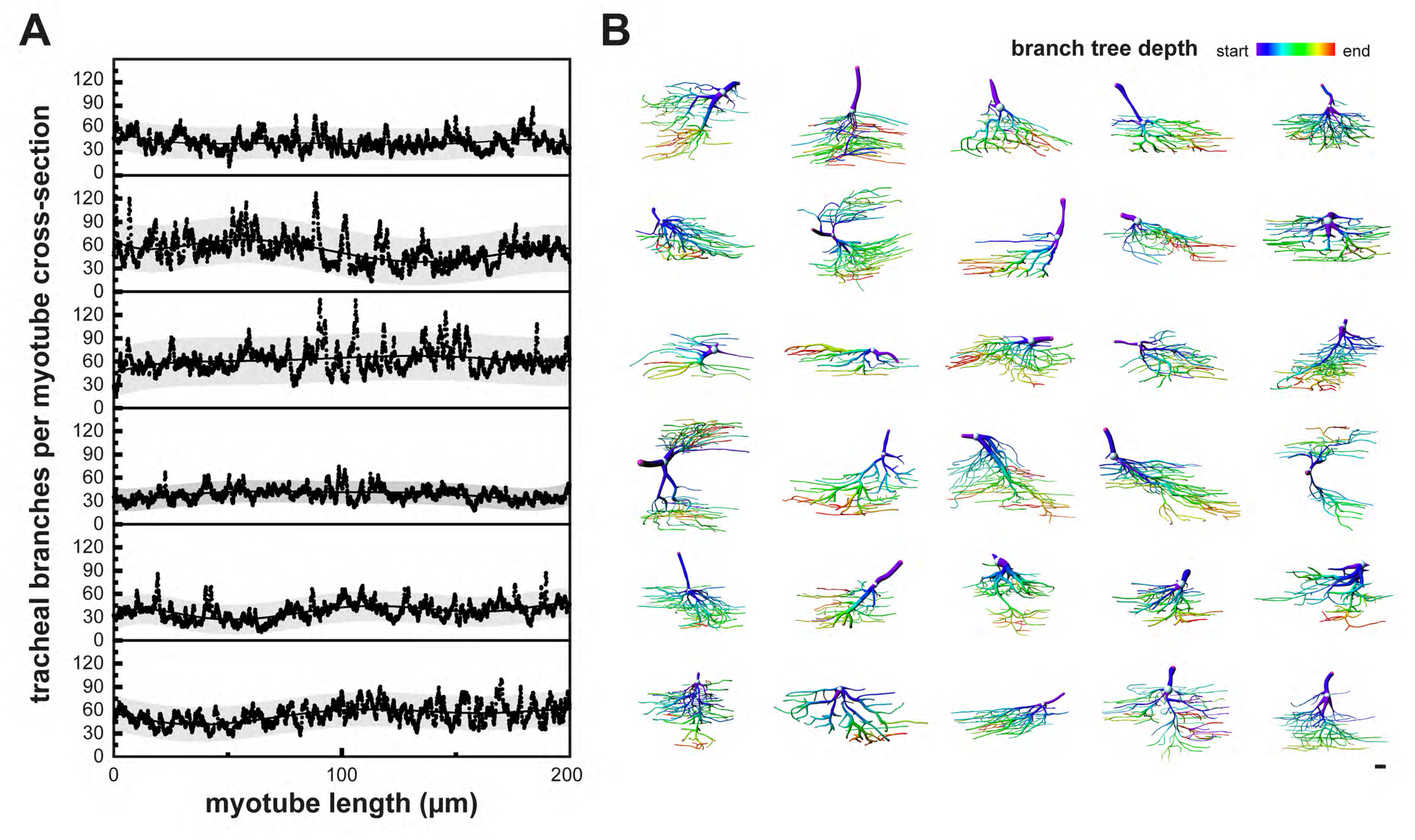
Tracheal terminal cells with non-stereotyped cellular morphologies fill the myotube volume. **(A)** The fraction of a myotube cross-section occupied by tracheal branches was plotted over a distance of 200 μm in 6 different myotubes. The grey band indicates the 95% prediction band. Note that myotubes are uniformly filled with tracheal branches along their length. **(B)** 30 individual segmented IFM tracheal terminal cells were color-coded for branch tree depth, which represents consecutive branching events. Warmer colors indicate higher-order branches. Note the heterogeneity in the tracheal cell morphologies. Scale bar: 10 μm (B).

**Supplementary Figure 2.**
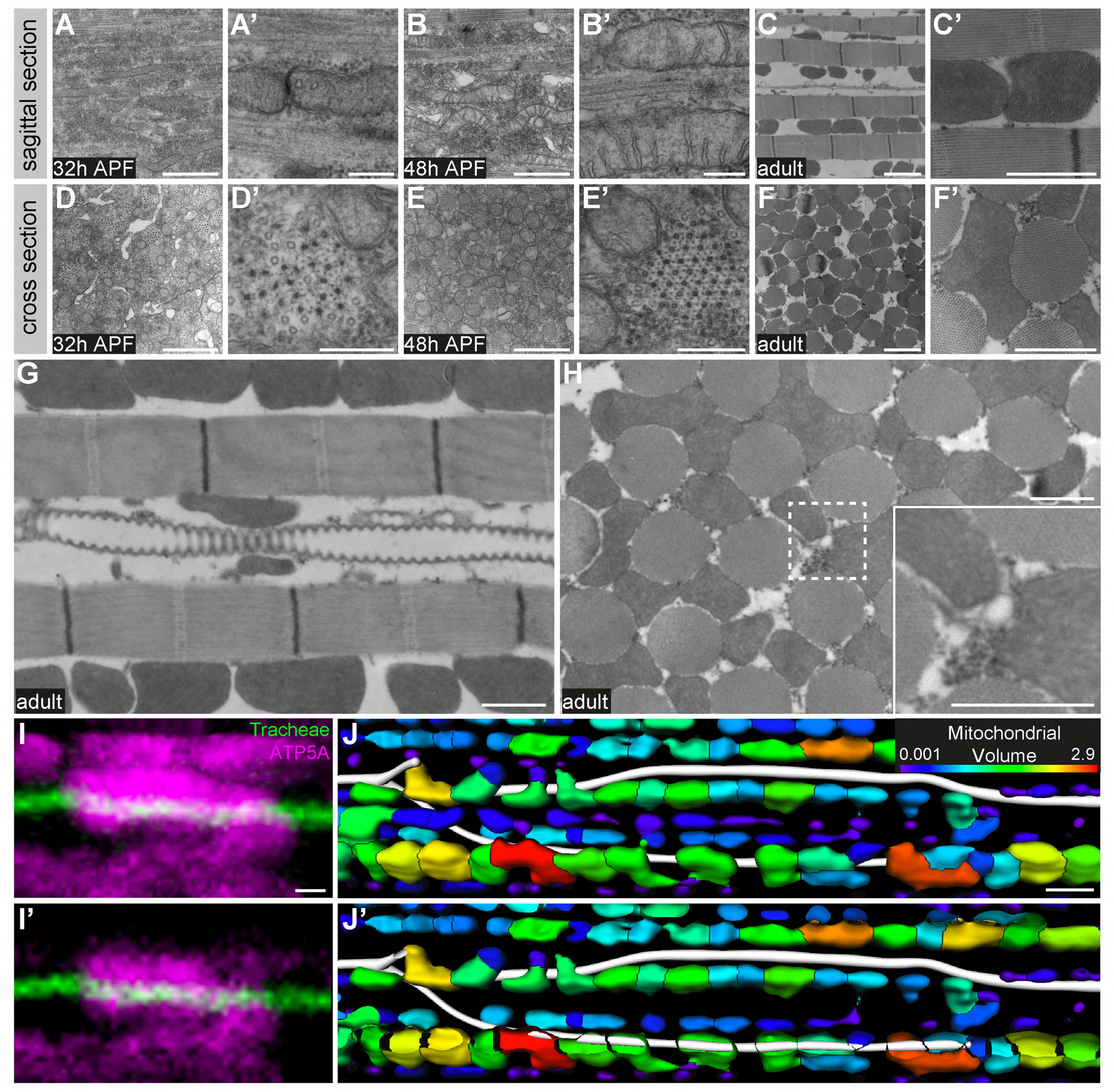
Mitochondria change their morphology during IFM development and partially enwrap tracheoles. **(A-H)** Transmission electron micrographs of sagittal **(A-C’,G)** and cross-sections **(D-F’,H)** of DLMs at 32 h APF **(A-D’)**, 48 h APF **(B-E’)** and in adult flies **(C-F’)**. Note that mitochondrial morphology changes from elongated-tubular (48 h APF) to globular shape in the adult. Cristae become increasingly elaborated as IFM development proceeds (compare **A’,B’** to **C’**). **(G,H)** Tracheoles are closely associated with mitochondria. The region marked by a dashed line in **H** is shown as a close-up (inset). **(I,I’)** show two consecutive confocal sections demonstrating partial enclosure of a tracheole (autofluorescence in green) by a mitochondrium labeled with anti-ATP5A antibody (magenta). Images were acquired using the Airyscan detector (LSM880) followed by Airyscan processing. **(J,J’)** Segmentation of tracheal branches (white) and mitochondria (color-coded for volume) in a myotube. **(J)** shows a projection of the entire z-stack, **(J’)** shows a projection starting from the plane with the tracheole. Note that mitochondria partially enwrap tracheoles. Scale bars: 1 μm (A,B,C’,D,E,F’,G,H), 200 nm (A’,B’D’,E’), 2 μm (C,F,J,J’), 0.4 μm (I,I’),.

**Supplementary Figure 3.**
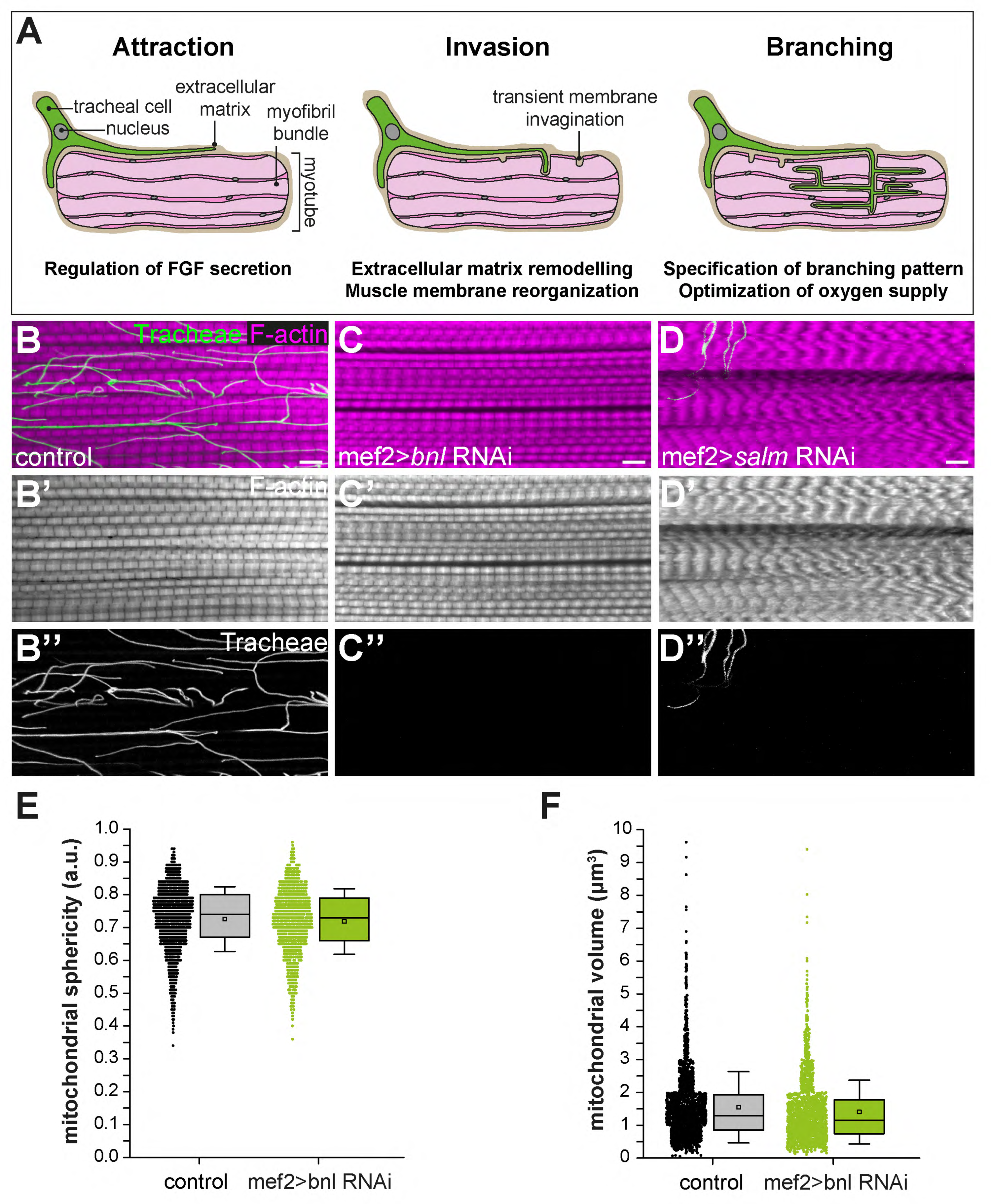
Identification of genes required for branch invasion. **(A)** Schematic illustration of the steps of IFM tracheation. Tracheal invasion into IFMs involves attraction by muscle-secreted Bnl FGF and entry of tracheal branches through membrane invaginations, which presumably provide access to the T-tubule system (Peterson & Krasnow, 2015). Ramification of tracheal branches within the myotube ensures efficient oxygen supply of mitochondria. **(B-D)** Close-ups of adult IFM myotubes of control flies (**B**), muscle-specific *bnl* RNAi (VDRC 109317; **C**) and muscle-specific *salm* RNAi (VDRC 3029; **D**). Myotubes were stained for F-actin (magenta), tracheae were visualized by their autofluorescence (green). Note that muscle-specific knock-down of *bnl* completely abolishes tracheal invasion but that myofibrils and sarcomeres of *bnl*-depleted trachealess IFMs do not show apparent morphological changes compared to wild-type controls (compare **B’** and **C’**). Further note that transformation of IFMs to tubular muscle fate upon depletion of *salm* (**D**) completely abrogates tracheal invasion. (**E,F**) Mitochondrial sphericity (**E**) and mitochondrial volume (**F**) are not altered in the tracheaIess IFMs of *bnl* tracheal knock-down-animals compared to wild-type controls. Scale bars: 5 μm (B-D).

**Supplementary Figure 4.**
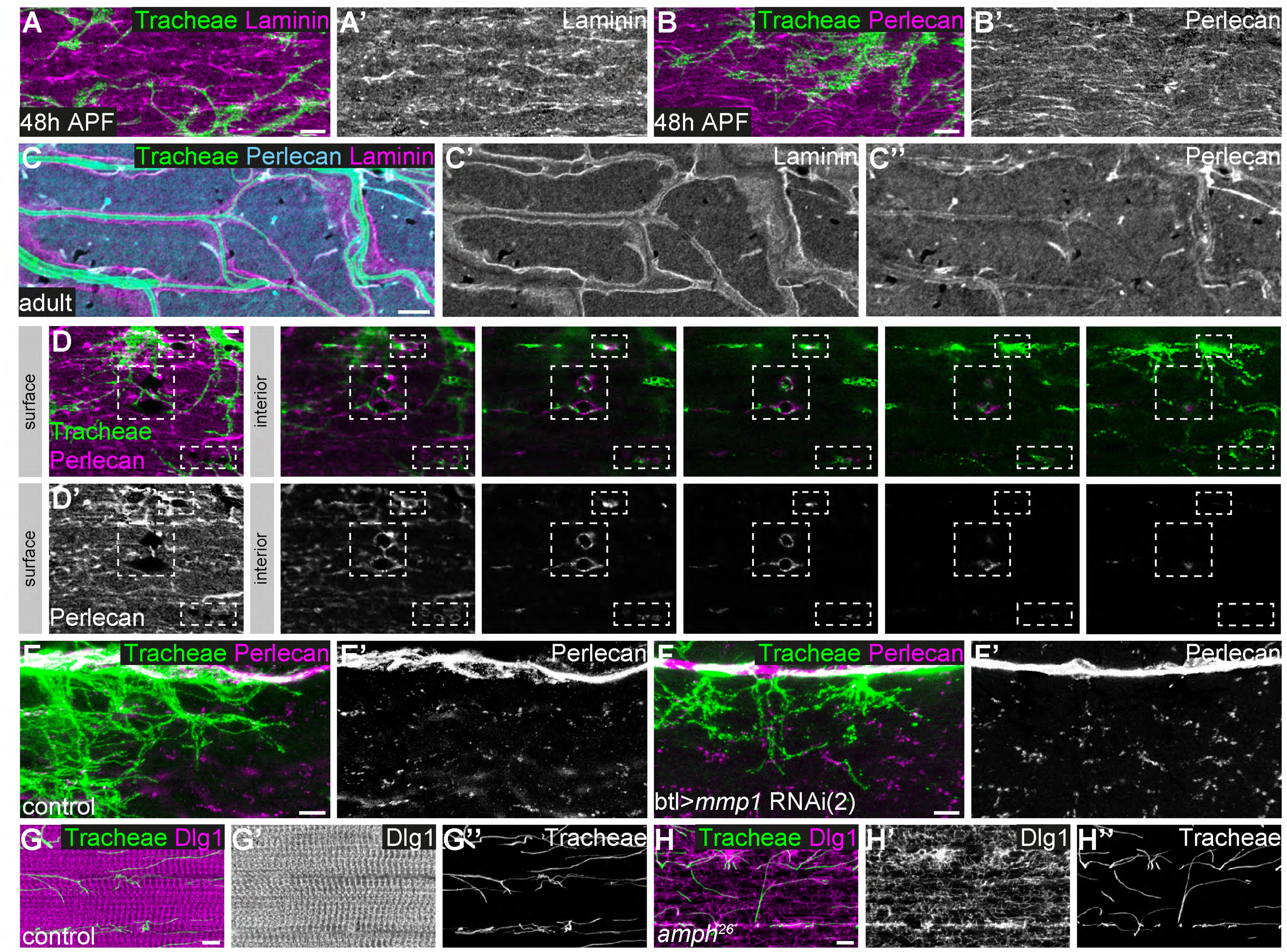
Basement membrane around tracheal branches increases during IFM development. **(A,B)** Localization of Laminin (magenta in **A**) and Perlecan (magenta in **B**) on the surface of developing IFMs 48 h APF. Tracheal cells express palmitoylated mKate2 (green) under the control of *btl*-Gal4. Note that at 48 h APF Laminin and Perlecan are mainly found on the myotube surface and not around invading tracheal branches. (**C**) In adult IFMs Laminin (magenta) and Perlecan (cyan) are found on the muscle surface and in basement membrane surrounding the tracheal branches (tracheal lumen visualized by autofluorescence; green). (**D**) Sequence of consecutive z-planes of a single myotube (48 h APF) from surface to interior. Perlecan (magenta) lines muscle membrane invaginations, which are being entered by tracheal branches marked by palmitoylated mKate2 (green). (**E,F**) Distribution of Perlecan around tracheal branches starting to invade IFMs 48 h APF in control (**E**) and tracheal *Mmp1* knock-down pupa (**F**). Note that the distribution of Perlecan around invading branches is not notably changed upon tracheal *Mmp1* knock-down. (**G,H**) Tracheal branching inside an adult myotube in control (**G**) and *amph*^*26*^ mutant (**H**). The T-tubule system is labeled by Dlg1 staining (magenta). Tracheal branches are visualized by their autofluorescence. Note that despite the severely disorganized T-tubule system in the *amph*^*26*^ mutant (Razzaq et al., 2001) (**H’**), tracheae have entered the myotube and show a normal branching pattern similar to controls (compare **H’’** to **G’’**). Scale bars: 5 μm (A-C’’, E-H’’) and 3 μm (D,D’).

## Supplementary Movies

**Supplementary Movie 1**

**Organization of tracheal branch invasion into a single IFM syncytium.**

3D animation of z-stack of a single DLM stained for F-actin (magenta). Immunostaining against LaminDm0 (cyan) labels all nuclei, DSRF (yellow) labels tracheal terminal cell nuclei. Tracheal branches (green) were visualized by their autofluorescence.

**Supplementary Movie 2**

**Mitochondria can enwrap IFM tracheoles.**

3D animation of an IFM tracheal branch surrounded by multiple mitochondria. Mitochondria were labeled by immunostaining against ATP5A (magenta). The tracheal branch (green) was visualized by its autofluorescence.

**Supplementary Movie 3**

**Dorsal view of tracheal invasion into DLMs in control and tracheal-specific *Mmp1* knock-down animals**

Time-lapse movies of tracheal invasion into DLMs in pupae 48 h APF. Tracheal cells are labeled by palmitoylated mKate2, IFMs are labeled by Myofilin-GFP. Dorsal views of a wild-type control pupa (top) and a tracheal *Mmp1* knock-down pupa (bottom) are shown. The movies were acquired with a 40x objective and a frame rate of 10 min in resonant scanning mode (Leica SP8) over 14 h. Z-stacks of ∼100 μm (0.35 μm step size) were imaged.

**Supplementary Movie 4**

**Lateral view of tracheal invasion into DVMs in control and tracheal-specific *Mmp1* knock-down animals**

Time-lapse movies of tracheal invasion into DVMs in pupae 48 h APF. Tracheal cells are labeled by palmitoylated mKate2, IFMs are labeled by Myofilin-GFP. Lateral views of a wild-type control pupa (top) and a tracheal *Mmp1* knock-down pupa (bottom) are shown. The movies were acquired with a 40x objective and a frame rate of 10 min in resonant scanning mode (Leica SP8) over 14 h. Z-stacks of ∼100 μm (0.35 μm step size) were imaged.

**Supplementary Movie 5**

**Close-up of invading tracheal branch tips in control and tracheal-specific *Mmp1* knock-down animals**

Close-up movies of tracheal invasion into DVMs in pupae 53 h APF. Tracheal cells are labeled by palmitoylated mKate2. A wild-type control (left) and a tracheal *Mmp1* knock-down pupa (right) are shown. Arrowheads point at growth cone-like structures. The movies were acquired with a 40x objective and a frame rate of 10 min in resonant scanning mode (Leica SP8). Z-stacks of ∼100 μm (0.35 μm step size) were imaged.

**Supplementary Movie 6**

**Distribution of ECM around single adult DLM with tracheae**

3D animation of ECM around a single adult DLM with tracheae. ECM components Laminin (magenta) and Perlecan (cyan) were visualized by immunostaining. Tracheal branches (green) were visualized by their autofluorescence. Note that tracheae enter the myotube through ECM-lined invaginations of the muscle surface.

## Supplementary Tables

**Supplementary Table 1**

**Quantitative analysis of IFM tracheal terminal cell morphology** Volume, sum of branch length, and total number of branch points and terminal points were extracted from 31 individually marked (MARCM clones) segmented terminal tracheal cells in wild-type pupae (75 h APF).

**Supplementary Table 2**

**RNAi screen to identify genes required for IFM tracheal invasion** Columns include gene identifiers and gene names of selected candidate genes, details about the RNAi lines used to knock down candidate genes, and the results of the screen for genes required for flight ability and tracheal invasion into myotubes.

